# Annotation-Based Gene-Peak Links Improve Regulatory Network Prediction of Gene Expression in Human Kidney Multi-Omics

**DOI:** 10.64898/2026.06.12.731741

**Authors:** Xinran Wang, Kimberly Siegmund, Jesse A. Goodrich, Jonathan Nelson, Huaiyu Mi, Lu Zhang, Steven Gazal, Bryan Queme, Darryl Shibata, Kelly Street

## Abstract

**Background:** Linking distal regulatory elements to their target genes is a central problem for interpreting chromatin accessibility and other non-coding genomic data. Proximity-based mapping is convenient but ignores three-dimensional enhancer-promoter architecture and can misassign long-range regulatory effects. Correlation-based approaches can also miss regulatory links because of limited statistical power and restrictive distance or significance thresholds. Single-cell and single-nucleus multi-omic datasets, such as 10x Multiome profiles that jointly measure chromatin accessibility and gene expression in the same cells or nuclei, now provide a way to evaluate gene-peak linkage strategies by testing how well linked accessibility features predict gene expression. Existing methods often focus on scoring individual enhancer-gene pairs. In this study, we proposed and constructed a fast annotation-based candidate gene-peak network and tested whether it improves downstream prediction of gene expression.

**Methods:** We first built a unified gene-peak regulatory network by integrating enhancer-based, promoter-based, and proximity-based linkage strategies. We then used single-cell multiome data from the Kidney Precision Medicine Project (KPMP) 10x Multiome cohort to evaluate whether the proposed links captured regulatory signals. We aggregated RNA expression and ATAC accessibility at the cell-type cluster level and trained predictive models to evaluate how well different linkage strategies could explain gene expression based on accessibility. Model performance was compared between annotation-based (including both enhancer- and promoter-based links) and proximity-based gene-peak links using testing R² and mean squared error (MSE) in a strict set of 1,704 genes and an adaptive set of 7,973 genes with more relaxed requirements.

**Results:** In the strict regime (1,704 genes with ≥20 peaks assigned by the nearest-TSS rule and ≥10 annotated enhancer-based peaks), the annotation-based model consistently achieved higher testing R² and lower testing MSE than the proximity-based model. In the larger adaptive regime (7,973 genes with ≥5 proximity-based and ≥2 enhancer-based peaks), we defined for each gene a balanced number of selected peaks based on its available closest and enhancer links; the annotation-based model again showed globally higher testing R² and lower testing MSE. These improvements were observed over a broad range of candidate-linked peak numbers.

**Conclusions:** Using human kidney 10x Multiome data, we show that a fast annotation-based gene-peak linkage framework can improve prediction of gene expression from chromatin accessibility compared with conventional approaches. These results support the use of biologically informed enhancer and promoter annotations when constructing candidate gene regulatory networks. Our framework also showed concordance with the correlation-based Signac LinkPeaks method while providing broader coverage and greater computational efficiency. We have implemented these annotation-based linkage methods in the GPlinksR R package, providing a fast and scalable tool for constructing regulatory networks.

## 1 Background

Linking distal regulatory elements to their target genes is a central problem in interpreting chromatin accessibility and other non-coding genomic data [1, 2]. Although it is common to assign each ATAC-seq peak to the nearest transcription start site, such proximity-based heuristics ignore the three-dimensional enhancer-promoter architecture of the genome and can misattribute regulatory effects, especially for long-range enhancers and genes with complex regulatory landscapes (Supplementary Figure 1) [3]. Functional and population-scale studies have shown that many enhancers skip the nearest gene, loop over hundreds of kilobases, and preferentially contact a subset of promoters within topologically associating domains (TADs), implying that traditional distance rules can both miss true connections and introduce spurious ones [1, 3]. In this context, we sought to develop a faster, comprehensive, and biologically informed strategy for linking genes and peaks more accurately than proximity-based mapping. To address this, we developed GPlinksR and then we used a single-cell 10x Multiome dataset to test whether the proposed gene-peak linkage network improved prediction of gene expression from chromatin accessibility compared with proximity-based links [4]. Correlation-based methods have also been proposed to link peaks and genes using joint accessibility-expression variation, but these approaches could miss regulatory links due to reliance on stringent distance or significance thresholds [5]. We therefore compared GPlinksR with Signac’s LinkPeaks method as a representative correlation-based approach.

The Kidney Precision Medicine Project (KPMP) [6] is a large NIH-funded prospective cohort study that collects clinically indicated kidney biopsies from adults, including healthy reference donors and patients with acute and chronic kidney disease across multiple U.S. centers. The study includes a healthy reference group and major disease groups such as chronic kidney disease (CKD) associated with diabetes or hypertension, acute kidney injury (AKI), and a diabetes-resilient cohort with long-standing diabetes but no clinical evidence of diabetic kidney disease (DM-R). KPMP generates deep molecular profiles from kidney tissue, including (i) an annotated single-nucleus RNA-seq (snRNA-seq) reference atlas and (ii) an initially unlabeled 10x Multiome dataset that measures paired RNA and ATAC profiles in the same nuclei. To enable downstream analysis of the Multiome data, we transferred cell-type labels from the snRNA-seq reference atlas to the Multiome RNA modality and then propagated the same labels to the matched Multiome ATAC modality using shared cell barcodes. This paired measurement of gene expression and chromatin accessibility across diverse kidney cell types makes KPMP well suited for benchmarking gene-peak linkage strategies. This analysis included 97 participants in total, comprising 40 healthy participants, 37 CKD, 17 AKI, and 3 DM-R participants. In this study, we used the KPMP 10x Multiome dataset as an evaluation dataset to test whether GPlinksR-derived annotation-based links improved gene expression prediction compared with proximity-based links, and to compare their overlap with correlation-based links from Signac LinkPeaks.

Several computational frameworks now address enhancer-gene linking, and they target different inferential goals. ABC and related extensions such as STARE score enhancer-gene pairs using enhancer activity and chromatin contact, with STARE adapting the ABC framework for broader single-cell applications. ENCODE-rE2G uses supervised modeling trained on perturbation-derived evidence to prioritize enhancer-gene links, and related work has used this framework to build large enhancer-gene resources. STREAM focuses on enhancer-driven regulatory network inference from paired single-cell transcriptome and chromatin accessibility data, and scMultiMap models enhancer-gene associations directly from sparse multimodal counts using a joint latent-variable framework [2,7–11]. These approaches are relevant to our problem, but they serve different inferential goals. Most aim to assign confidence to individual enhancer-gene pairs or infer regulatory structure from activity, correlation, or perturbation evidence. GPlinksR is designed to provide a complementary framework for rapidly assembling a broad candidate gene-peak regulatory network that can serve as a scalable backbone for downstream prediction and modeling, rather than a replacement for pair-specific statistical or perturbation-based methods.

In this study, we leveraged the GPlinksR framework, which combines three linkage sources: enhancer-based links (AnnoQ/PEREGRINE [12–14]), promoter-based links (EnsDb.Hsapiens.v86 [15]), and proximity-based links (nearest TSS) to construct an integrated gene-peak regulatory network for the KPMP human kidney 10x Multiome dataset. To evaluate whether the proposed links improved prediction of gene expression from chromatin accessibility, random forest models [16] were trained to predict cluster-averaged gene expression from linked peak accessibility. By comparing proximity-only models with models that also incorporated enhancer and promoter information, we quantified the added predictive value of annotation-based links. This design allowed us to show GPlinksR’s framework performance. Our goal was not to compete with existing methods on per-pair classification or statistical inference. Instead, we constructed a fast, comprehensive candidate regulatory network for capturing biologically informed gene-peak linkages.

## 2 Methods

### 2.1 Data preprocessing (Figure 1)

Single-cell RNA-seq and paired single-cell ATAC-seq from 10x Multiome experiments were obtained from the KPMP consortium [6]. Because RNA and ATAC measurements arose from the same physical nuclei, all downstream analyses were performed on cells with matched RNA and ATAC profiles. A previously processed KPMP snRNA-seq atlas with fine-grained annotations (subclass.l2) was used as the reference for cell label transfer.

**Figure 1.**
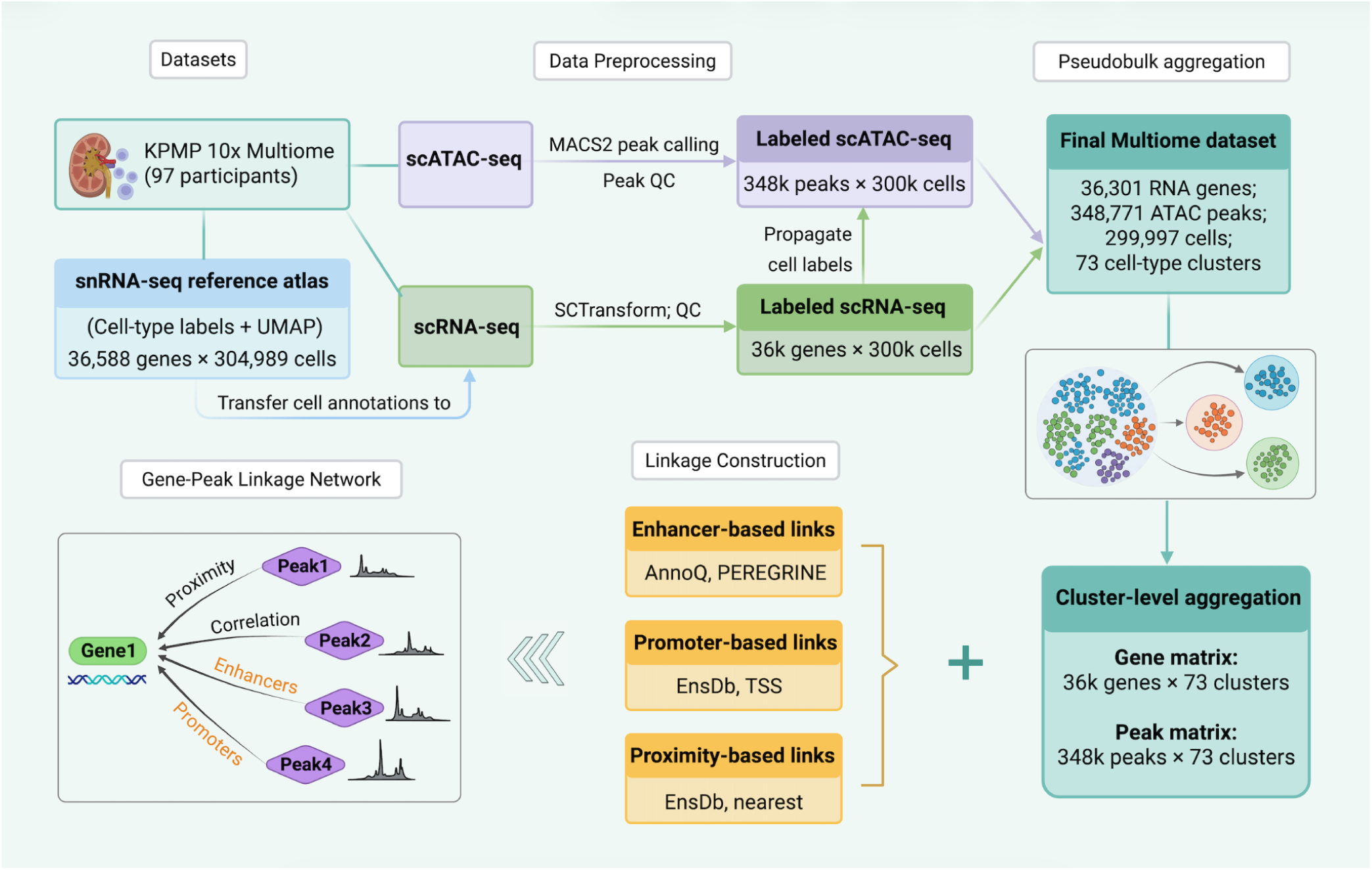
Overview of the multiome data processing and gene-peak linkage construction workflow [17]

For RNA, counts matrices from 97 participants were read from 10x Genomics HDF5 files and converted into Seurat objects. Cells were identified using a combination of participant identifiers and cell barcodes. Quality control removed low-quality or likely doublet cells, including cells with fewer than 200 or more than 10,000 expressed genes and cells with mitochondrial read fraction greater than 5%. Following QC, participant-level objects were merged and normalized using SCTransform [18]. Principal component analysis (PCA) and UMAP were performed using the standard Seurat workflow. The resulting merged RNA object contained 392,170 cells and 36,301 genes.

To transfer fine-grained annotations from the KPMP snRNA-seq reference (36,588 genes and 304,989 cells), the merged RNA object was treated as the query dataset. The reference data was processed in parallel. Highly variable genes were defined as the intersection of variable feature sets from the reference and query, yielding approximately 1,300 shared genes. Transfer anchors were identified using FindTransferAnchors with normalization.method set to “SCT” and reduction set to “pcaproject”, using 30 principal components from the reference. Predicted cell identities and associated prediction scores were obtained with TransferData. MapQuery was then used to project the query cells onto the reference UMAP for visualization and cross-dataset comparison. The transferred labels were stored in the RNA object as the “predicted.id” identity class for downstream aggregation.

For ATAC, fragment files from all 97 participants were processed with Signac. Fragment coordinates were combined into a pooled BED file and subjected to MACS2 [19] peak calling against the hg38 reference genome, producing a narrowPeak set. Peaks were filtered to retain high-confidence events, requiring signalValue greater than 1.3 and qValue greater than 2 (false discovery rate less than 0.01). Filtered peaks were converted to GRanges, and gene annotations were obtained from EnsDb.Hsapiens.v86 using UCSC-style chromosome naming. For each participant, a count matrix was constructed using FeatureMatrix over the unified peak set.

Primary ATAC quality control retained cells with total peak counts between 100 and 20,000, nucleosome signal less than 4, and TSS enrichment greater than 2. Sample-level objects were prefixed with participant identifiers and merged in batches into a unified ATAC object (348,771 peaks, 344,230 cells). For single-cell analysis, we extracted count matrices from processed Seurat objects and restricted analyses to barcodes present in both datasets. To avoid exceeding the sparse-matrix size limit (2^31−1) during sample-level merge operations, we applied uniform global downsampling (proportion = 0.87) of the barcodes after overlap filtering, so that all retained cells were sampled using the same proportion. The final constructed single-cell dataset contained 299,997 shared cells, yielding an RNA matrix of 36,301 genes × 299,997 cells and an ATAC matrix of 348,771 peaks × 299,997 cells. Cell-type annotations were taken from the “predicted.id” labels in the RNA object and used to define 73 distinct cell-type clusters. For downstream modeling, cells assigned to the same annotated cluster were pooled across participants.

For each gene in the scRNA-seq data, we summarized gene expression at the cell-type cluster level. Specifically, within each of the 73 annotated cell-type clusters, cells were randomly split into 70% training and 30% testing subsets, and normalized gene expression was averaged separately within the training and testing cells. This produced, for each gene, a 73-dimensional training response vector and a corresponding 73-dimensional testing response vector, where each entry represents the mean expression of that gene within one cluster. We used an analogous aggregation strategy for chromatin accessibility at linked peaks.

Conceptually, this approach resembles cell type-level pseudobulk aggregation. We adopted this strategy because gene expression and chromatin accessibility exhibit consistent patterns within cell types, and cluster-level aggregation reduces the sparsity inherent to single-cell data. Directly modeling gene counts at the single-cell level is challenging due to extreme sparsity and high dimensionality (36,301 genes × 299,997 cells). This aggregation reduces noise and yields a more stable and computationally tractable prediction framework. Features with missing values, constant values, or near-zero variance were removed prior to modeling.

### 2.2 Regulatory gene-peak linkage construction

We constructed a unified regulatory network by combining three complementary types of gene-peak links: enhancer-based links, promoter-based links, and proximity-based links. After standardizing chromosome names across data sources, peak-gene links from the three sources were merged and deduplicated.

Enhancer-based links were derived from the PEREGRINE hg38 enhancer resource [12] as distributed through the AnnoQ platform [13], which integrates PANTHER-based regulatory annotations [14]. The PEREGRINE resource integrates enhancer annotations from ENCODE, Ensembl, FANTOM, and VISTA, together with multiple experimental evidence types (e.g., Hi-C, ChIA-PET, eQTL), providing broader coverage and more comprehensive enhancer-gene linkage information [12]. Enhancer records were read as genomic intervals and mapped to HGNC gene identifiers, which were then converted to gene symbols using Ensembl biomaRt. Only genes present in the RNA assay were retained. Enhancer intervals were converted to genomic ranges, and chromosome names were standardized to match the ATAC peak set. Overlaps between enhancers and ATAC peaks were computed and unique gene-peak pairs were retained. This procedure yielded 231,476 enhancer-based gene-peak links (Figure 2). The distribution of enhancer-based linked peaks per gene is shown in Supplementary Figure 2.

**Figure 2.**
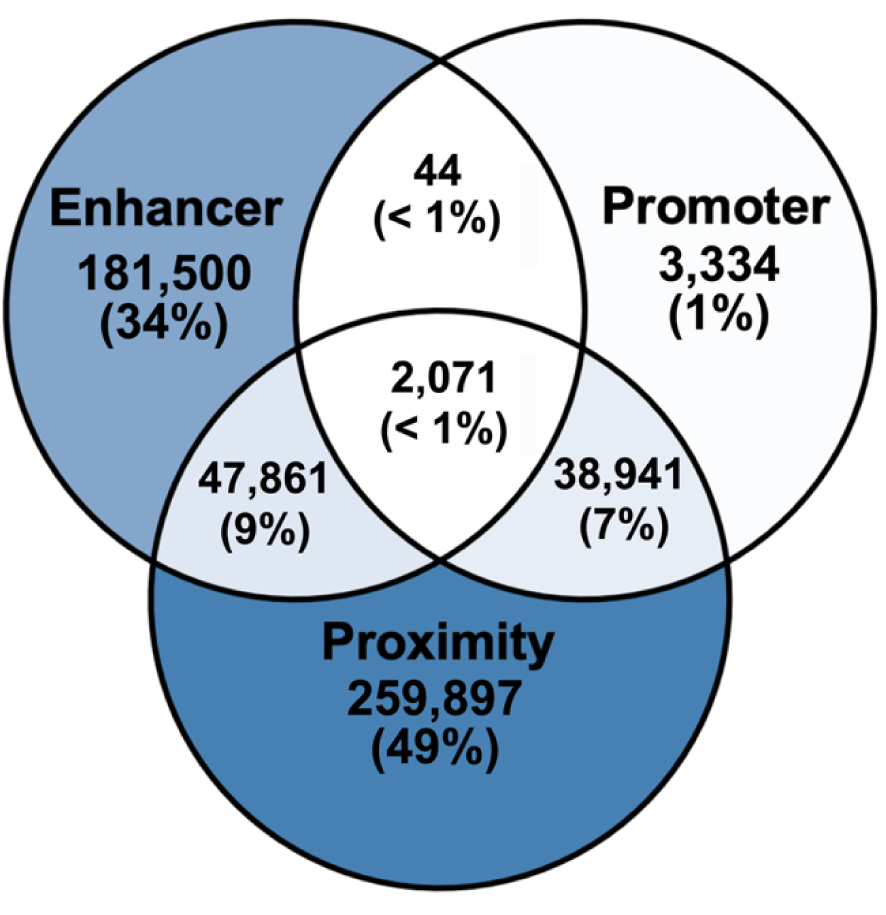
Composition and overlap of enhancer-, promoter-, and proximity-based gene-peak links in the KPMP regulatory network

Promoter-based links were defined using transcription start sites (TSS) from EnsDb.Hsapiens.v86 [15]. For each gene, we constructed promoter windows spanning 2,000 base pairs upstream and 200 base pairs downstream of the TSS. After harmonizing seqlevels with the ATAC peaks, we overlapped promoter windows with the unified peak set. For each overlapping pair, we recorded a promoter-based link between the gene and the peak, again keeping one record per unique gene-peak pair. This produced 44,390 promoter-based gene-peak links. As shown in Figure 2, only 3,334 of these links were promoter-specific; 38,941 overlapped with proximity-based links, 44 overlapped with enhancer-based links, and 2,071 were shared across all three linkage strategies.

Proximity-based links were constructed by assigning each ATAC peak to its single nearest transcription start site (TSS). TSS coordinates were defined from EnsDb.Hsapiens.v86, and distances were measured between the center of the peak and the TSS. For each peak, the gene corresponding to the nearest TSS was identified using the Bioconductor nearest function on genomic ranges. The resulting proximity-based network comprised 348,771 gene-peak links (Figure 2). The distribution of proximity-based linked peaks per gene is shown in Supplementary Figure 3.

All three link sets were harmonized by ensuring consistent chromosome naming and peak identifiers across sources. Duplicated entries across sources were removed, and the final link table contained, for each gene-peak pair, the peak coordinate, the gene symbol, and a source label indicating enhancer, promoter, closest, or combinations thereof if the same pair appeared in multiple sources. After merging and deduplication, the unified network included 22,035 genes, 348,771 unique ATAC peaks, and 533,648 unique gene-peak links. The distribution of total gene-peak linkage counts per gene in the integrated Gene-Peak link network is shown in Supplementary Figure 4. This network served as the basis for feature selection in the predictive models. For each gene, we computed cluster-level mean chromatin accessibility for the selected predictor peaks and used these values to predict the corresponding 73-dimensional cluster-level gene-expression profile.

### 2.3 Machine-learning framework for evaluating gene-peak links

To evaluate gene-peak linkage strategies, we used a machine-learning framework in which cluster-level gene expression was predicted from chromatin accessibility at linked peaks. The unit of analysis was the annotated cell-type cluster. Within each of the 73 clusters, cells were randomly split into 70% training and 30% testing subsets, and RNA and ATAC signals were aggregated separately within each split. Thus, for each gene, models were trained on 73 aggregated training observations and evaluated on 73 aggregated testing observations, with one observation per cluster. Two feature sets were defined for each gene: Model 1 used proximity-based peaks assigned via the nearest-TSS rule, and Model 2 extended these features by incorporating annotation-based links. Random forest regression models (randomForest package) were fitted after excluding genes with zero variance or missing values and removing predictors with missing or constant values. Random forest was selected here as a flexible and robust model that captures nonlinear relationships and interactions with relatively limited tuning. Because our goal was to compare gene-peak linkage strategies rather than optimize predictive performance, we used a single consistent modeling framework to ensure that observed differences reflect the linkage strategy rather than model selection or tuning.

We defined two regimes that differ in gene and peak selection to ensure a fair comparison between Model 1 and Model 2. This design isolates the contribution of annotation-based features from differences in feature count:

In the strict regime, designed to evaluate performance under abundant regulatory information, we restricted analysis to genes with at least 20 closest peaks and 10 enhancer peaks. Model 1 used the 20 most variable proximity-based peaks. Model 2 used 10 proximity-based and 10 enhancer-based peaks, ranked by variability. Promoter-linked peaks were included in both models when available. Because promoter links largely overlapped with proximity links (38,941 of 44,390; Figure 2) and contributed relatively few unique pairs (3,334), their incremental effect was expected to be limited. Hyperparameters were tuned within the training data using repeated 3-fold cross-validation. Each candidate model was fit using 150 trees. The initial number of variables considered at each split was set to min(7, p), where p is the number of predictors, and tuning considered the number of variables per split, minimum node size, and maximum tree size. Final models were refitted using 500 trees on the full training data and evaluated on the held-out test set. Matching the number of selected peaks across models allowed us to isolate the effect of feature type. This regime included 1,704 genes.

In the adaptive regime, designed to reflect more realistic data conditions, we relaxed the criteria to require at least five closest peaks and at least two enhancer peaks to ensure a stable feature set, resulting in 7,973 genes. For each gene, we counted the number of available candidate closest peaks, *n_C_* and candidate enhancer peaks, *n_E_* and defined a balanced number of selected peaks as *x* = *min*([*n_C_*/2], *n_E_*). This choice ensured that both models used a comparable total number of peaks and preserved a one-to-one balance between closest and enhancer features in Model 2. Model 1 used the top 2*x* closest peaks ranked by variability, whereas Model 2 used the top *x* closest peaks and top *x* enhancer peaks ranked by variability; all available promoter peaks were also added to both models as stated in the strict regime. This design maintains comparable model size and balances feature sources. Random forest models were fitted as above, with the number of variables considered at each split initialized as ceiling(*p*/3) and bounded between 1 and p, and tuned using repeated cross-validation within the training data. Final models were refitted with 500 trees and evaluated on the test set.

For each gene, we recorded the total number of candidate-linked peaks available before selection (*n_peaks_*) and the resulting *x* value. By controlling the number of predictors across models, performance differences primarily reflect the contribution of annotation-based features rather than model size. Model performance was summarized using the testing coefficient of determination (R²) and testing mean squared error (MSE). For each gene, we computed R² between predicted and observed cluster-level expression in the test set and the corresponding MSE. Genes with constant predictions or zero variance in test expression were excluded from R² summaries but retained for MSE. To visualize how performance varied with the number of candidate linked peaks per gene (i.e., the number of unique linked peaks per gene across closest, enhancer, and promoter sources), we plotted individual gene-level metrics against this count and overlaid generalized additive model (GAM) [20] smooth curves.

### 2.4 Software and implementation

All analyses were performed in R version 4.4 or higher. Data handling and single-cell processing were conducted using Seurat, Signac, SeuratDisk, and Matrix, with genome annotation facilitated by biomaRt, GenomeInfoDb, and EnsDb.Hsapiens.v86. Peak calling was carried out using MACS2 [19]. Regulatory networks and gene-peak link construction were implemented in the GPlinksR package (https://github.com/Corawang123/GPlinksR), which provides functions for assembling enhancer, promoter, and proximity links from external resources and for exporting unified gene-peak tables suitable for downstream modeling. Random forest models were fitted using the randomForest package, and visualization of performance metrics used ggplot2. All computations were executed in a high-performance computing environment.

## 3 Results

### 3.1 Proximity-only models provide a baseline for expression prediction

Cluster-level aggregation produced RNA and ATAC matrices comprising 73 cell-type clusters, approximately 36,000 genes, and 349,000 peaks. For genes in the strict regime, proximity-only models (Model 1) predicted cluster-level gene expression using a fixed set of 20 selected closest peaks per gene, yielding a broad spectrum of testing R² values (Figure 3A). In Figure 3A, the x-axis represents the number of candidate linked peaks available for that gene before selection. Testing performance tended to improve as the candidate pool increased.

**Figure 3.**
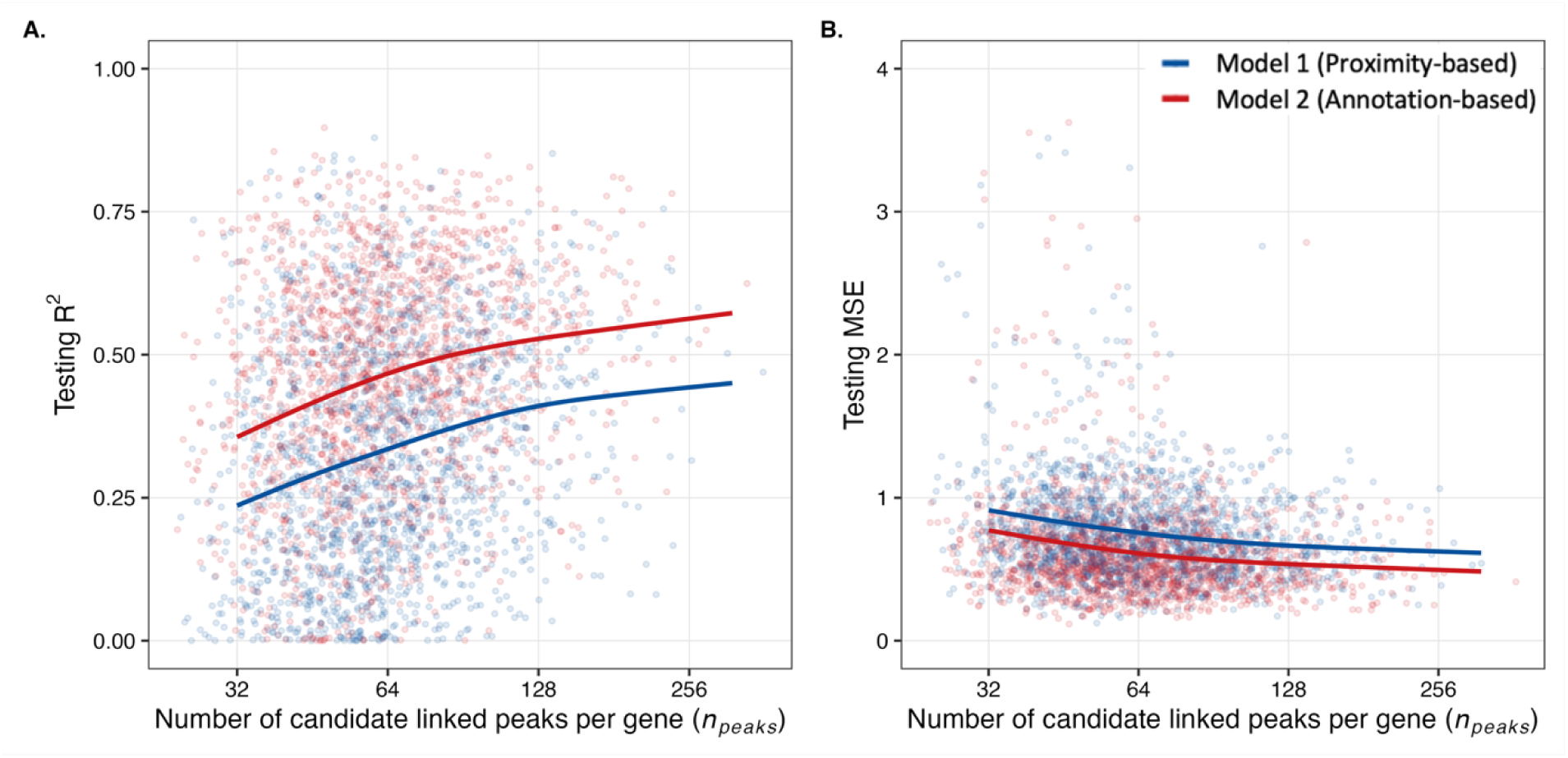
Performance comparison between proximity-based (Model 1) and annotation-based (Model 2) models in the strict regime as a function of the number of candidate linked peaks per gene. (A) Testing R² for predicting cluster-level gene expression. (B) Testing MSE for predicting cluster-level gene expression. Points show per-gene performance; solid lines show smoothed trend curves across genes.

### 3.2 Enhancer-based models improve prediction in the strict gene set

When enhancer peaks were added in the strict regime, Model 2 substantially outperformed the proximity-only Model 1. In Figure 3A, the GAM smooth for testing R² is consistently shifted upward for Model 2 across the full range of candidate linked peak counts, indicating that the benefit of enhancer-based features is not restricted to genes with especially large candidate peak pools. This separation suggests that enhancer peaks contribute additional, non-redundant regulatory information beyond what is captured by nearby peaks alone.

The same conclusion was supported by testing MSE. In Figure 3B, the GAM smooth for Model 2 remained below that of Model 1 across the entire range of candidate linked peak counts, indicating lower prediction error. A paired Wilcoxon signed-rank test across the 1,704 genes in the strict regime confirmed this pattern: Model 2 achieved higher testing R² than Model 1 (median 0.474 vs 0.319; P < 0.0001) and lower testing MSE (median 0.562 vs 0.708; P < 0.0001). Model 2 improved R² in 1,520 of 1,704 genes (89.2%) and reduced MSE in 1,531 of 1,704 genes (89.8%). Together, these results show that adding enhancer peaks materially improves gene-expression prediction in the strict gene set.

### 3.3 Adaptive, balanced peak selection extends improvements to 7,973 genes

To determine whether the advantage of annotation-based features extends beyond the densely linked strict gene set, we analyzed an adaptive set of 7,973 genes that met the relaxed requirement of at least five closest peaks and at least two enhancer peaks. For each gene, we defined a balanced number of selected peaks. This design mirrors a more realistic setting, in which genes differ widely in the number of linked peaks while still allowing Model 1 and Model 2 to be compared using the same total numbers of features.

In Figure 4A, Model 2 shows higher testing R² across most of the observed range of candidate linked peak counts, with the GAM smooth remaining above that of Model 1 from genes with relatively small candidate peak sets to those with larger candidate pools. Figure 4B shows the complementary pattern for testing MSE, where the Model 2 smooth remains below that of Model 1 across nearly the full range of candidate linked peak counts. These patterns indicate that the benefit of adding enhancer-based features is maintained across a broad spectrum of genes rather than being confined to a narrow subset with unusually dense linkage structure.

**Figure 4.**
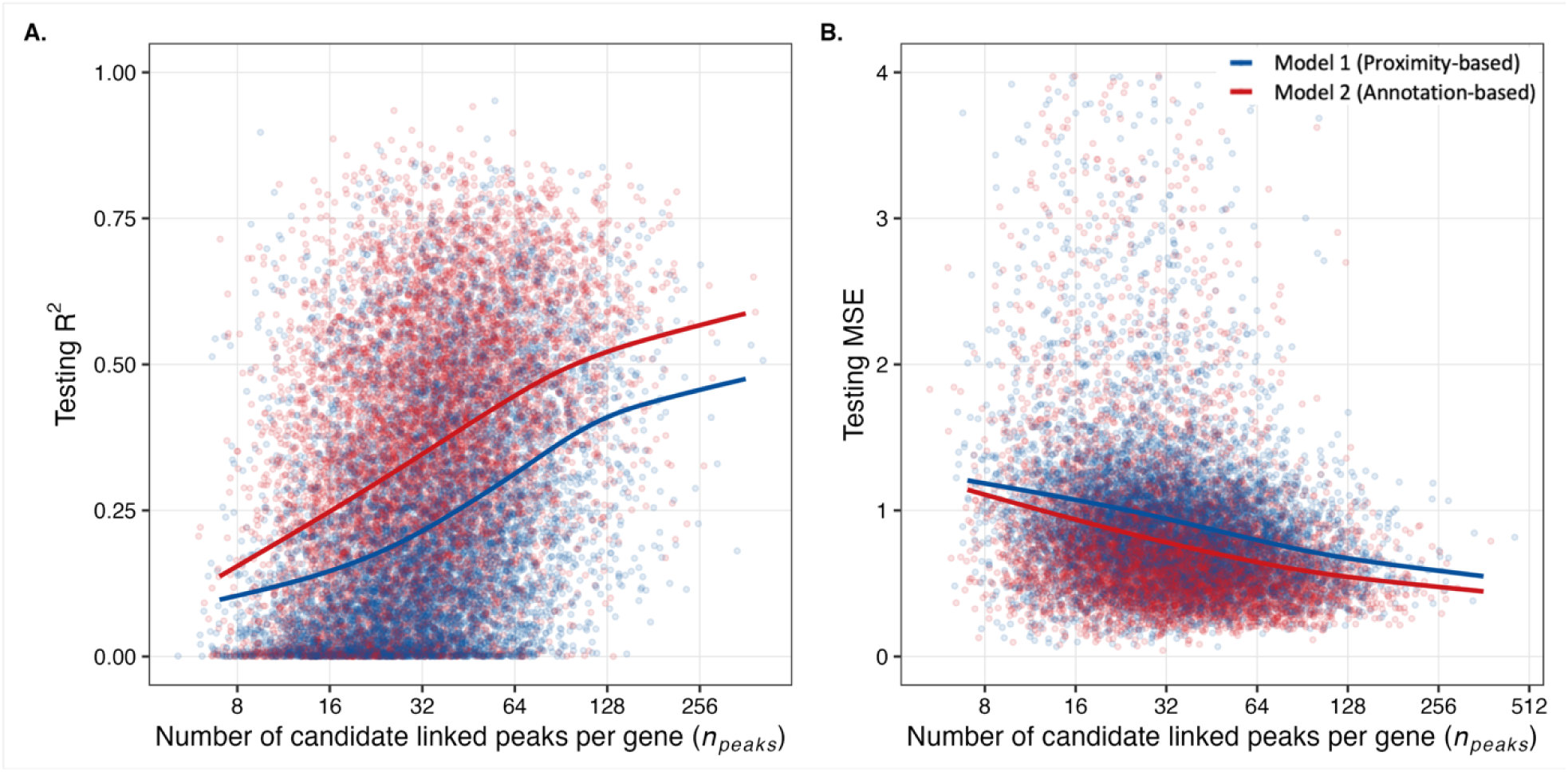
Performance comparison between proximity-based (Model 1) and annotation-based (Model 2) models in the adaptive regime as a function of the number of candidate linked peaks per gene. (A) Testing R² for predicting cluster-level gene expression. (B) Testing MSE for predicting cluster-level gene expression. Points show per-gene performance; solid lines show smoothed trend curves across genes.

The paired gene-level comparison supported the same conclusion. Across 7,973 genes, Model 2 achieved higher testing R² than Model 1 (median 0.350 vs 0.180, paired Wilcoxon signed-rank test, P < 0.0001) and lower testing MSE (median 0.675 vs 0.850, paired Wilcoxon signed-rank test, P < 0.0001). Model 2 improved R² for 6,663 genes (83.6%) and reduced MSE for 6,643 genes (83.3%). Together, these results indicate that enhancer-based peak selection remains associated with better predictive performance under the adaptive design, even in a larger and more heterogeneous gene set.

### 3.4 Consistency between strict and adaptive regimes

The strict and adaptive regimes differ in gene selection and feature composition. However, they lead to the same qualitative conclusion. When Model 1 and Model 2 were matched for the number of selected peaks per gene, models incorporating enhancer- and promoter-based links outperformed models based on proximity-based links. In the strict regime, the improvement was most pronounced for genes with dense regulatory annotation. These genes had many enhancer peaks, and those peaks were strongly informative. In the adaptive regime, the effect was more moderate but remained consistent across thousands of genes and across a wide range of candidate peak counts.

The agreement between the two regimes suggests that the improvement is not driven by a specific feature-selection rule or a restricted gene subset. Instead, it reflects a general property of enhancer and promoter annotations. When combined with chromatin accessibility data, these annotations help identify regulatory peaks that carry predictive information about gene expression.

We also examined an additional factor, gene-expression variance, that may influence prediction performance. Supplementary Figures 5 and 6 show that genes with higher variance across clusters were generally easier to predict. These genes showed both higher testing R² and larger gains from Model 2. This suggests that prediction performance is also related to gene-expression variance. These results are consistent with the main findings. Annotation-based links improve prediction by expanding the set of candidate regulatory features beyond closest peaks. This effect is stronger for genes with complex or distal regulatory structure. Together, these findings support the motivation and use of biologically informed linkage strategies to predict gene expression from chromatin accessibility.

### 3.5 Comparison with Signac LinkPeaks()

Building on the strict/adaptive model comparisons above, we further benchmarked our gene-peak link framework against Signac’s correlation-based LinkPeaks() method [21], a commonly used approach for identifying putative regulatory interactions in single-cell multi-omic (scATAC-seq + scRNA-seq) datasets. Signac’s LinkPeaks function identifies gene-peak associations by testing Pearson correlations between peak accessibility and gene expression for peaks within 500 kb of each gene’s transcription start site (TSS), while applying internal quality control steps that require sufficient signal, variability, and activity across cells. To ensure a standard implementation, we followed the default settings recommended in the Signac LinkPeaks() documentation, including requiring a peak or gene to be detected in at least 10 cells, using 200 background samples, and retaining links with correlation score and p-value cutoffs of 0.05. This design produces a conservative set of highly active gene-peak links, but it also substantially restricts the number of genes and peaks that are ultimately tested and reported.

When applied to the KPMP multiome dataset (36,301 genes, 348,771 peaks), LinkPeaks produced 5,247 unique gene-peak links, involving only 3,197 genes and 4,927 ATAC peaks. In contrast, our annotation-based GPlinks framework yielded a much more comprehensive regulatory network comprising 533,648 gene-peak links, including 22,035 genes and 348,771 peaks. Importantly, annotation-based gene-peak links contributed 273,751 unique links, accounting for 51.30% of all links in the regulatory network (Figure 2).

Despite this difference in scale, pairwise comparison still revealed substantial agreement between the two approaches. Of the 5,247 unique gene-peak pairs identified by Signac, 4,590 links overlapped exactly with our previously constructed gene-peak regulatory network (Figure 5). Thus, 87.48% of Signac-derived links were represented in our gene-peak link set. Among these 4,590 overlapping links, 4,191 (91.31%) were proximity-based, 241 (5.25%) were enhancer-based, and 158 (3.44%) were promoter-based. These results show that the GPlinks network captures most links identified by Signac’s LinkPeaks while extending substantially beyond them in scale and coverage.

**Figure 5.**
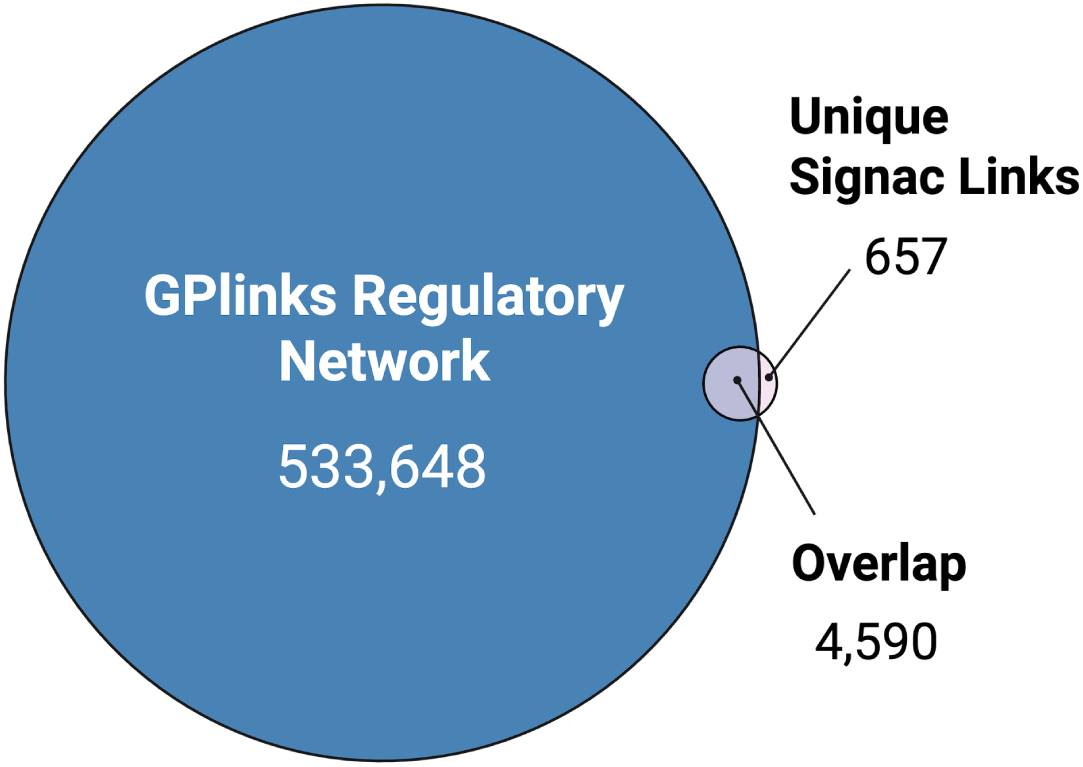
Overlap of gene-peak links identified by annotation-based GPlinks approach and Signac LinkPeaks function

The remaining 657 Signac-only links represent correlated peak-gene pairs that are not captured by the current GPlinks network. These links highlight a known limitation of annotation-based approaches: incomplete or context-specific regulatory annotations. As such, these cases motivate future expansion of enhancer and promoter resources and illustrate how correlation-based methods can complement annotation-driven networks by flagging candidate regulatory interactions that lack prior annotation support.

### 3.6 Computational efficiency and scalability

Beyond differences in coverage, GPlinks and LinkPeaks differ substantially in computational cost. In our implementation, running Signac’s LinkPeaks across all genes and all peaks required approximately 1 day and 6 hours of wall-clock time on an HPC R environment, reflecting the heavy computational burden of genome-wide correlation testing and empirical null estimation over large numbers of peak-gene pairs. In contrast, the GPlinks gene-peak link set can be constructed from enhancer, promoter, and proximity annotations in under 30 minutes when implemented through the GPlinksR package. It does not require GPU resources or high-core-count compute and can be run on a local computer with sufficient RAM. The GPlinks framework enables rapid construction of regulatory networks across tissues, cohorts, or parameter settings, making it well suited for large-scale studies, exploratory analyses, and iterative downstream modeling workflows that would be difficult or impractical to perform using correlation-based methods alone.

## 4 Discussion

In this study, we used single-cell multi-omic data from the human kidney to evaluate how different gene-peak linkage strategies influence the ability to predict gene expression from chromatin accessibility. We first compared (i) proximity-only models and (ii) annotation-based models that integrate enhancer-, promoter-, and proximity-based links constructed using our GPlinks framework. We then benchmarked two classes of gene-peak links: (i) annotation- and proximity-based link sets generated by the GPlinks framework and (ii) correlation-based links identified using Signac’s LinkPeaks() method. Together, these analyses allowed us to assess both the predictive utility and the broad coverage of annotation-based gene-peak linkage strategies.

### 4.1 Predictive modeling benefits of Gene-Peak regulatory network

The predictive modeling results highlight the value of annotation-based gene-peak networks. Across genes, random forest models trained using GPlinks-derived regulatory features consistently achieved higher predictive accuracy than models based on proximity-based peak sets alone. This improvement suggests that incorporating external enhancer and promoter annotations allows the model to capture regulatory elements that are not necessarily the closest peaks to a gene but still contribute to gene expression. In particular, genes with more complex regulatory networks, reflected by a larger number of linked peaks, appear to benefit from this richer candidate feature space. This finding emphasizes the importance of annotation-based linkage strategies when constructing regulatory networks for downstream predictive and deep learning analyses.

### 4.2 Relation to existing enhancer-gene linking methods

Our results should be interpreted in the context of existing enhancer-gene linking methods and are not intended as a replacement. Methods such as ABC/STARE and ENCODE-rE2G focus primarily on prioritizing enhancer-gene pairs, STREAM and scMultiMap address broader network inference or association modeling in single-cell multimodal data [7–11]. The GPlinksR framework is designed for a different use case. It assembles an annotation-based candidate gene-peak network and then evaluates whether these candidate links improve prediction of gene expression from chromatin accessibility. Its main strengths are coverage, computational efficiency, and ease of integration into downstream modeling workflows, rather than per-pair precision.

We also emphasize that GPlinksR builds on substantial prior work in this field. The enhancer-gene annotations used through AnnoQ/PEREGRINE draw on enhancer resources such as ENCODE, FANTOM, Ensembl, and VISTA [22–25], as well as studies that developed or evaluated enhancer-gene linking frameworks, including ABC, STARE, ENCODE-rE2G,

STREAM, and scMultiMap [7–11], among others. We are grateful to the authors of these studies and resources, as our framework benefits from their data, methods, and accumulated biological insight. GPlinksR is therefore best viewed as a downstream integrative application of existing biological knowledge, not as a replacement for the methods that made such analyses possible.

This allows the annotation-based GPlinks network to recover many of the highly expressed, highly accessible genes targeted by correlation-based methods, and also to incorporate lowly expressed or transcriptionally silent genes whose regulation may still be biologically meaningful but difficult to detect through correlation alone.

This difference in purpose is also important when interpreting the comparison between GPlinks and Signac’s LinkPeaks. Signac’s LinkPeaks is correlation-based and intentionally conservative, returning links that only show detectable correlation in the observed data. As a result, it primarily captures regulatory relationships for highly active genes and highly accessible regions, while excluding genes with low expression, low accessibility, or limited variability across cells. In contrast, GPlinks is designed to construct a biologically informed candidate regulatory network that emphasizes biological plausibility and broader coverage over statistical correlation significance. This allows the annotation-based GPlinks network to recover many of the highly expressed, highly accessible genes targeted by correlation-based methods, and also to incorporate lowly expressed or transcriptionally silent genes whose regulation may still be biologically meaningful but difficult to detect through correlation alone.

### 4.3 Limitations and future usage

Several limitations of the present work should be noted. First, the GPlinks network depends on the quality and completeness of external enhancer and promoter annotations, which remain imperfect and tissue dependent. Second, our evaluation relies on cluster-level aggregation, which simplifies cell-to-cell heterogeneity and may obscure transient or state-specific regulation. Third, we treat all linked peaks equally once selected, without explicitly modeling differences between enhancer-, promoter-, and proximity-based interactions.

We also did not benchmark GPlinksR against CRISPRi ground truth or directly compare it with methods such as ABC/STARE, ENCODE-rE2G, STREAM, or scMultiMap on a shared pairwise evaluation task [7–11]. This reflects both scope and data availability: GPlinksR constructs an annotation-based candidate network rather than directly estimating pair-specific enhancer-gene association scores or probabilities, and, to our knowledge, kidney-specific perturbation resources are not yet available at a scale that would support a matched benchmark in this setting. Future work should revisit these comparisons as larger perturbation-based kidney datasets become available.

In addition, because training and testing cells were split within clusters rather than by participant, cells from the same participant could contribute to both sets, which may make absolute prediction performance somewhat optimistic. The strict regime focuses on genes with a relatively large number of linked peaks (≥20 proximity-based and ≥10 enhancer-based links), representing a non-random subset enriched for genes with more complex or active regulatory architectures; therefore, results from this subset may not fully generalize to genes with sparser regulatory landscapes. These genes may also be more likely to exhibit enhancer-mediated regulation and higher expression variability, which may partly explain the observed performance gains.

Despite these limitations, the GPlinks framework provides a foundation that can be extended as annotation resources improve and as more sophisticated modeling approaches become available. An important strength of the GPlinks network is its suitability for downstream machine learning and deep learning applications. By providing a dense, biologically informed regulatory graph spanning both active and inactive genes, it may serve as a structured prior for models that aim to learn regulatory logic, cellular plasticity, or context-dependent regulatory programs from multi-omic data.

## 5 Conclusions

Linking chromatin accessibility to gene expression remains challenging, as proximity-based rules often overlook long-range regulatory interactions, and correlation-based approaches are usually computationally intensive and tend to recover associations among highly transcriptionally active genes. Using matched single-cell RNA and ATAC data from the Kidney Precision Medicine Project, we show that integrating enhancer and promoter annotations with proximity information provides a scalable and biologically grounded gene-peak linkage network. Regulatory networks constructed with this framework consistently improve gene expression prediction relative to proximity-based links across a wide range of genes and regulatory contexts.

Beyond predictive performance, a major strength of this approach lies in its coverage and efficiency. The GPlinks regulatory network recovers most links identified by correlation-based methods such as Signac’s LinkPeaks while substantially expanding regulatory coverage to additional genes and regulatory regions, including those with lower activity. Because this network can be constructed rapidly without genome-wide correlation testing, it provides a practical foundation for large-scale analyses. Rather than replacing pairwise statistical or perturbation-based methods, GPlinks may serve as a complementary and scalable regulatory backbone for downstream modeling and for studies of gene regulation activity across cell types and conditions.

## Acknowledgements

The Kidney Precision Medicine Project (KPMP) is supported by the National Institute of Diabetes and Digestive and Kidney Diseases (NIDDK) through the following grants: U01DK133081, U01DK133091, U01DK133092, U01DK133093, U01DK133095, U01DK133097, U01DK114866, U01DK114908, U01DK133090, U01DK133113, U01DK133766, U01DK133768, U01DK114907, U01DK114920, U01DK114923, U01DK114933, U24DK114886, UH3DK114926, UH3DK114861, UH3DK114915, and UH3DK114937. We gratefully acknowledge the essential contributions of our patient participants and the support of the American public through their tax dollars.

---

**Supplementary Figure 1.**
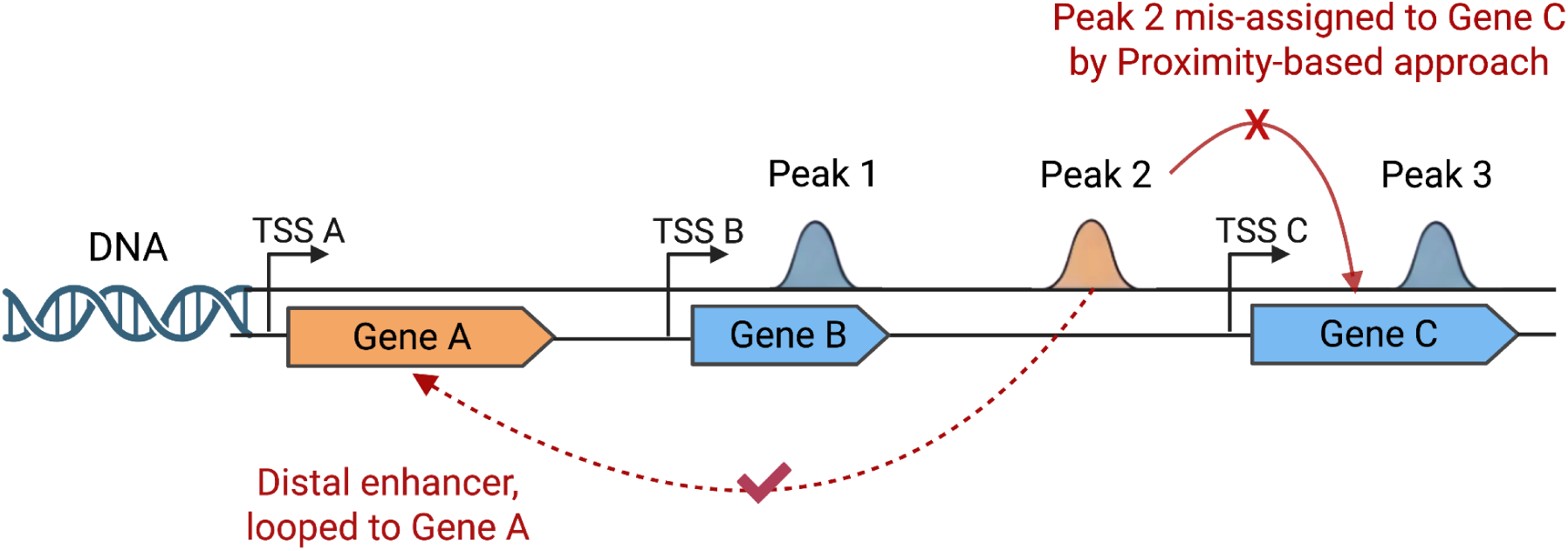
Illustration of distal enhancer-gene interactions and potential misassignment by proximity-based peak-gene linkage

**Supplementary Figure 2.**
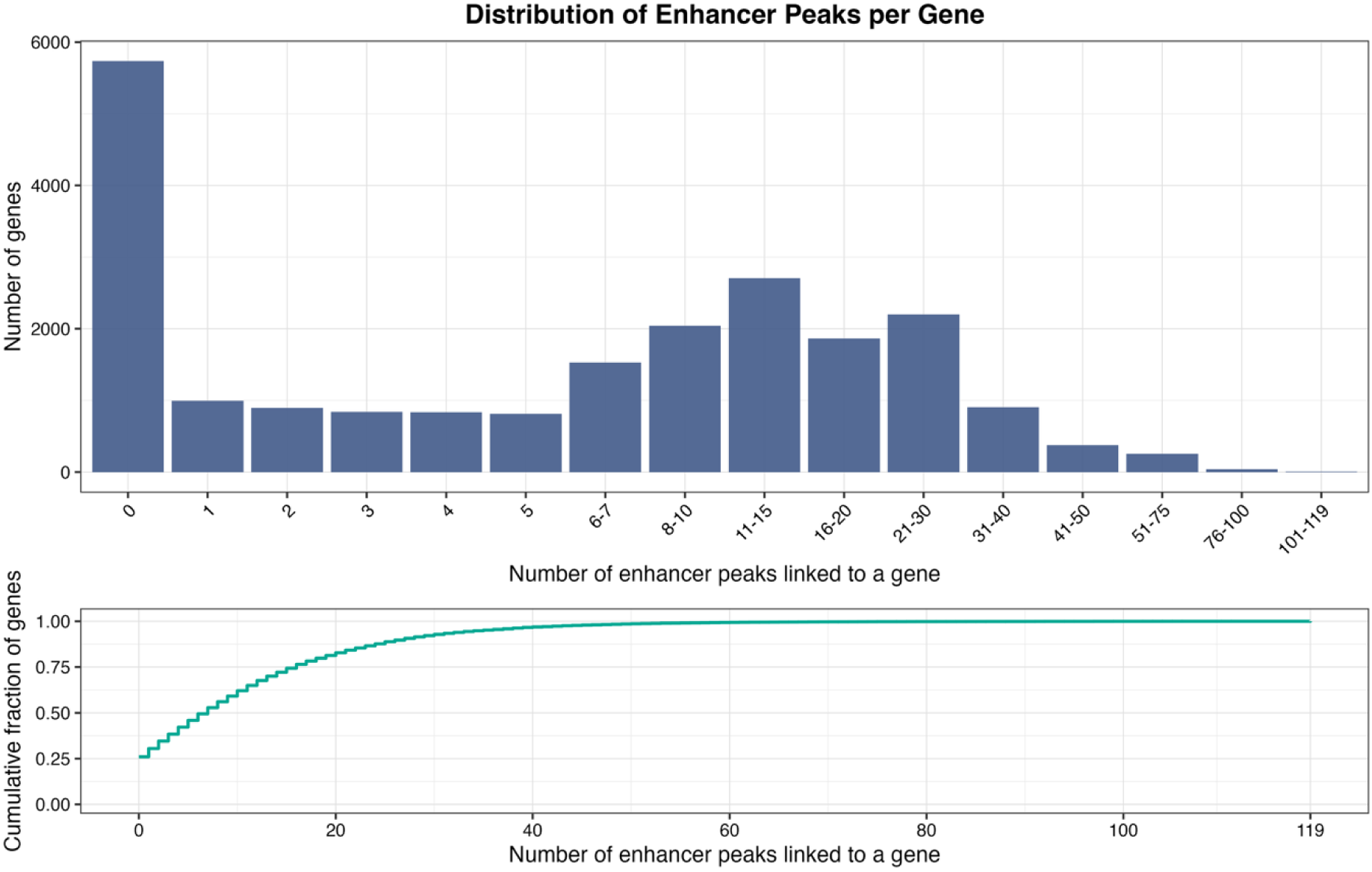
Distribution of enhancer-based linked peaks per gene across all 22,035 genes

**Supplementary Figure 3.**
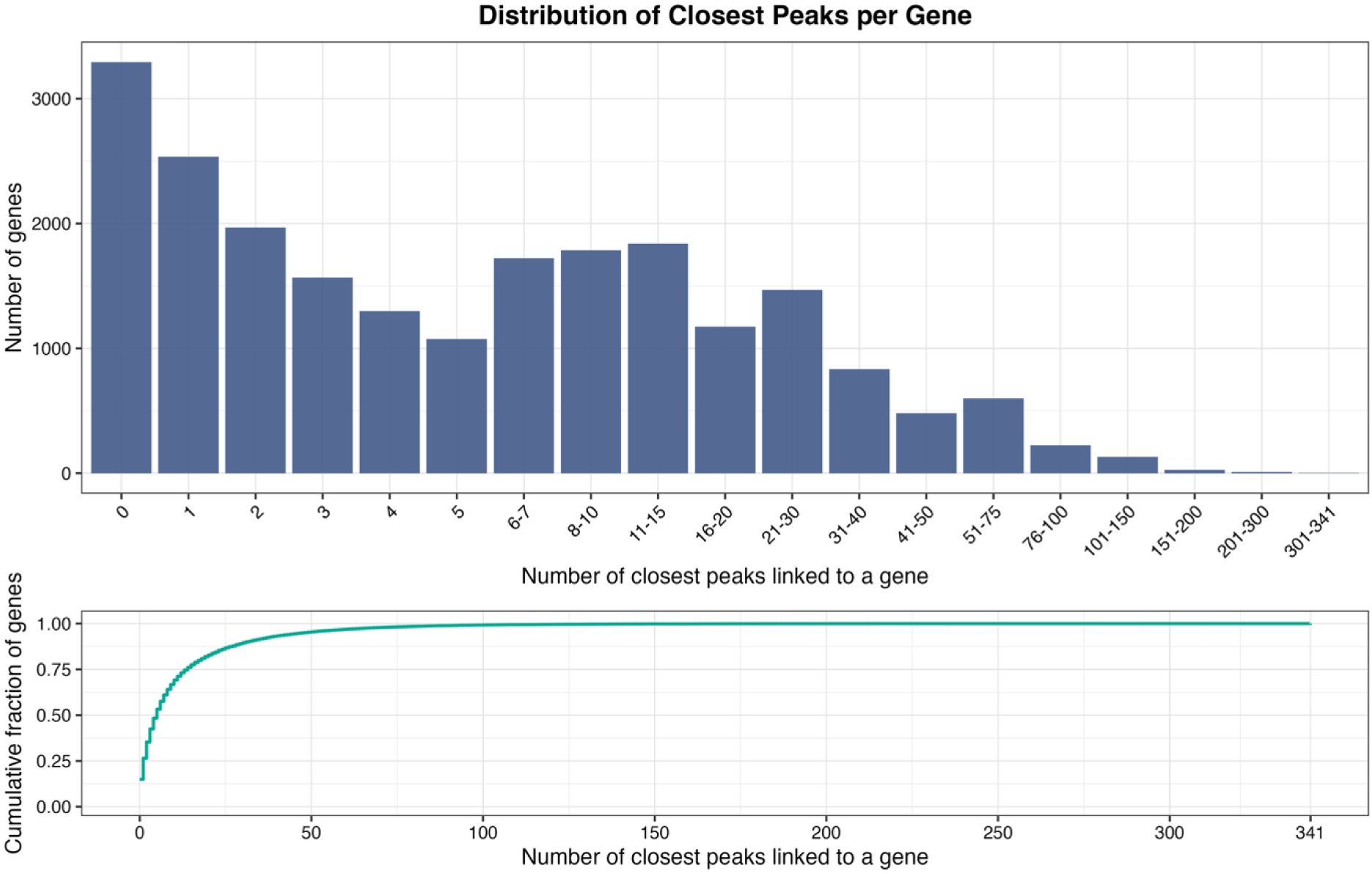
Distribution of proximity-based linked peaks per gene across all 22,035 genes

**Supplementary Figure 4.**
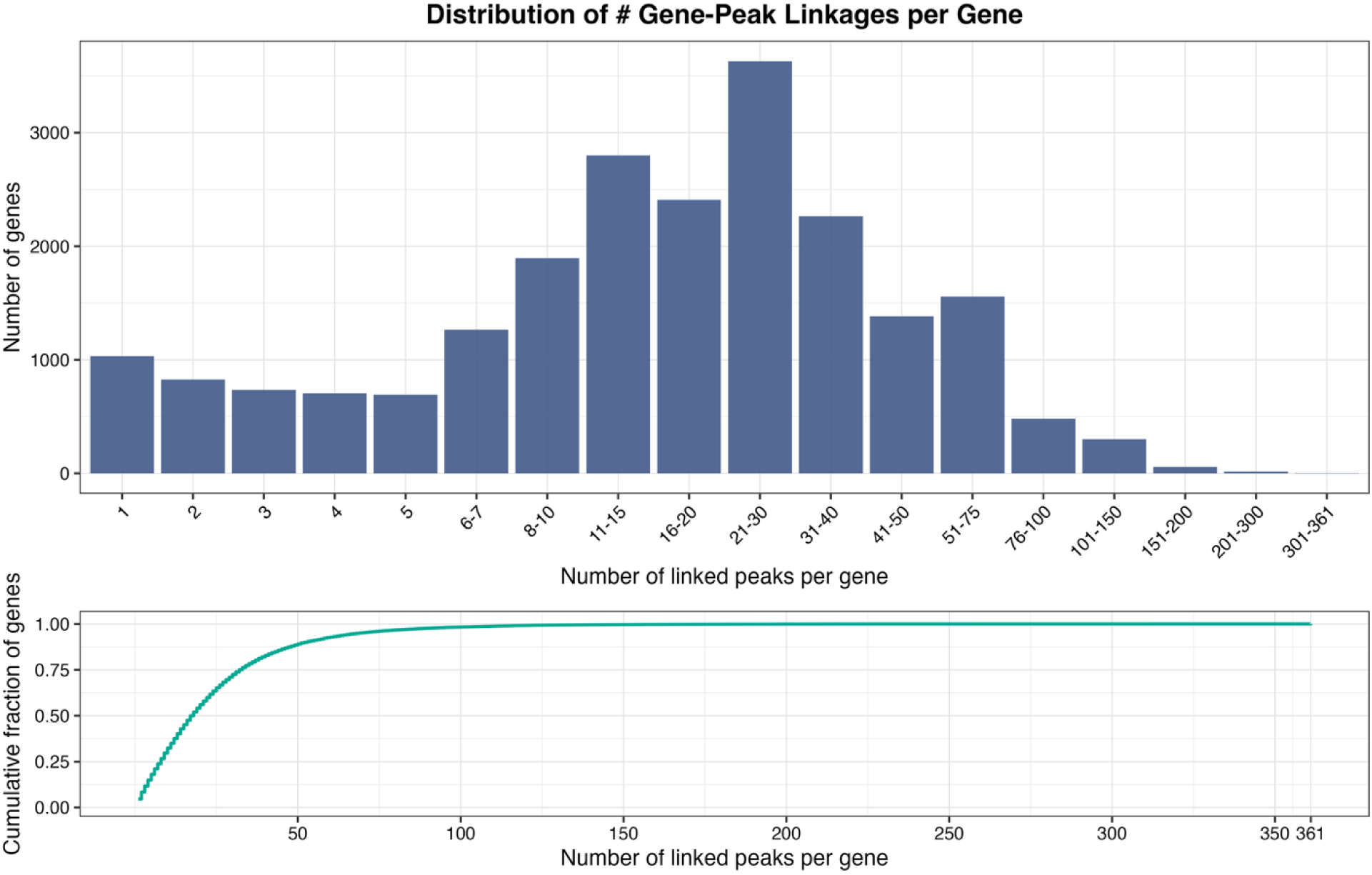
Distribution of gene-peak linkage counts per gene.

**Supplementary Figure 5.**
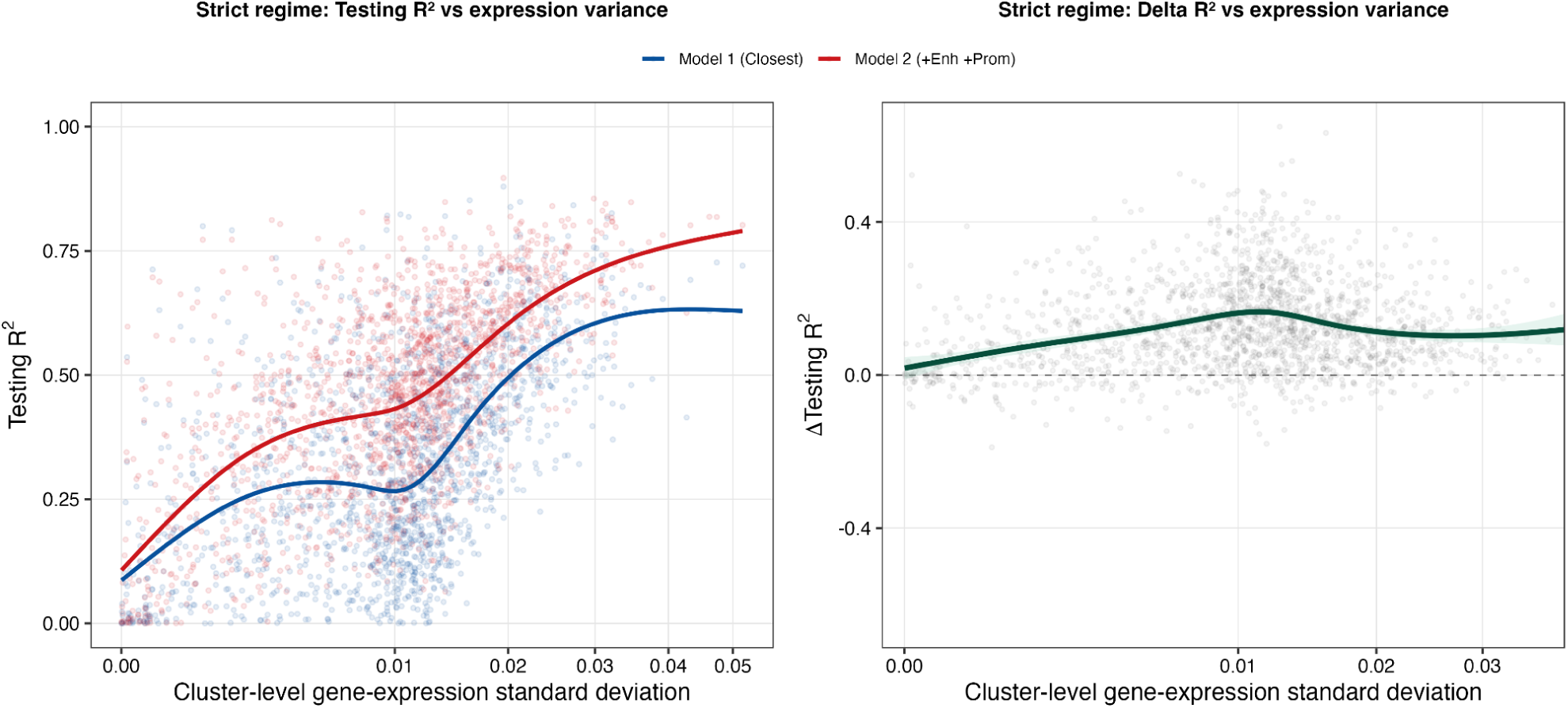
Association of model prediction performance with cluster-level gene-expression variance in the strict regime. Left panel: testing R² for Model 1 and Model 2 as a function of cluster-level gene-expression variance. Right panel: per-gene improvement in predictive performance (ΔR² = R² of Model 2 - R² of Model 1) as a function of cluster-level gene-expression variance.

**Supplementary Figure 6.**
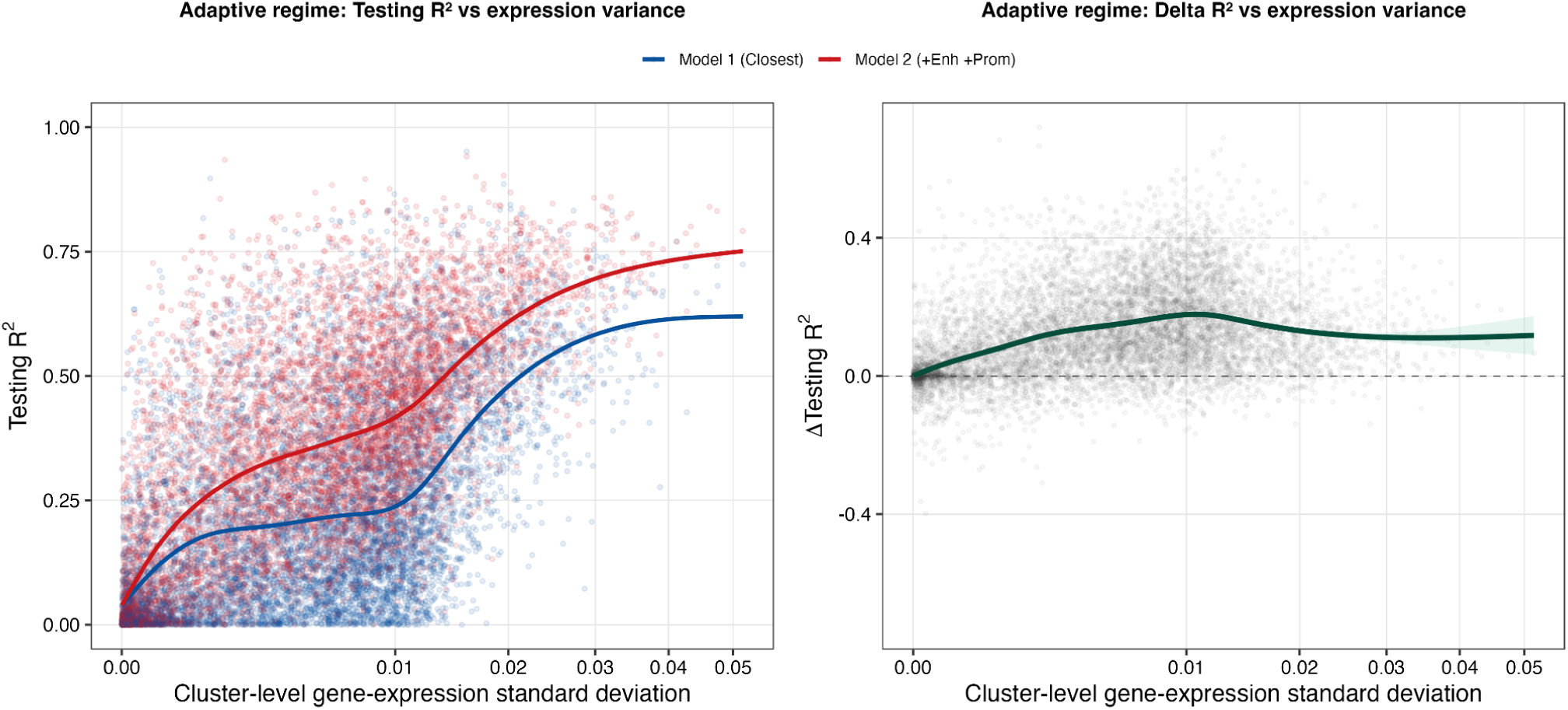
Association of model prediction performance with cluster-level gene-expression variance in the adaptive regime. Left panel: testing R² for Model 1 and Model 2 as a function of cluster-level gene-expression variance. Right panel: per-gene improvement in predictive performance (ΔR² = R² of Model 2 - R² of Model 1) as a function of cluster-level gene-expression variance.

